# REAP supplemental fertilizer improves greenhouse crop yield

**DOI:** 10.1101/2020.08.27.266916

**Authors:** Rachel Backer, Damian Solomon

**Author notes:** Corresponding author: Rachel Backer, 21,111 Lakeshore Rd, Sainte-Anne-de-Bellevue, Qc, H9X 3V9, Canada.

## Abstract

**Background:** Mine tailings contain rare earth elements, including lanthanum and cerium, and plant micronutrients including iron. Previous studies have demonstrated that fertilizers containing rare earth elements and/or micronutrients can influence plant physiology, nutrient uptake and crop yield. However, applying the right dose of these fertilizers is critical since the concentration range associated with benefits is often narrow, and overapplication can lead to crop yield reductions. This study aimed to quantify the effects of a water-soluble fertilizer, REAP, on the yield of greenhouse crops.

**Methods:** In the first experiment, the effects of three concentrations of REAP (100, 250 or 500 ppm) were compared to a control (0 ppm REAP) on growth of lettuce, tomato and pepper growing in soilless media. In the second experiments, the effects of REAP applied at higher rates (500, 1000 and 2000 ppm) were compared to a control (0 ppm REAP) on the growth of lettuce, peppers, tomato and cantaloupe.

**Results:** In the first experiment, there were no significant differences in yield between treatments, REAP appeared to promote root development. In the second experiment, there were significant yield increases for all crops treated with REAP. Gas exchange rates and nutrient concentration of tomato plants receiving REAP were not significantly different from the control. These results demonstrated that nutrient elements in REAP, including lanthanum, cerium, and micronutrients, improved the growth and yield of vegetable crops when applied at rates ranging from 500 to 2000 ppm.

## Introduction

Plant nutrition and fertilization is a balance between preventing nutrient deficiencies, due to under-fertilization, or toxicity, due to overfertilization, while achieving economic returns resulting from increased crop yields. In hydroponic greenhouse crop cultivation, nutrients are supplied as liquid fertilizers with balanced nutrient compositions that meet crop macro- and micro-nutrient requirements. However, the electrical conductivity of these solutions must be carefully monitored to avoid causing osmotic stress in crops. One strategy for stimulating crop growth while reducing osmotic stress in hydroponic greenhouse crops is application of fertilizers containing lanthanide elements, such as lanthanum and cerium (Haneklaus et al., 2015) and micronutrients including iron.

Research undertaken in China has suggested that soil application of these elements has the potential to improve crop yields. Mine tailings, leftover after the extraction of zinc, are a rich source of these nutrients. Now that mine tailings elements are becoming more readily available in the US, these products can be developed at a price that is viable for the production of greenhouse or controlled environment horticultural crops, including lettuce, tomatoes, peppers, cantaloupe and cannabis.

### Rare earth elements effects on plant physiology

Lanthanides are rare earth elements present in plant tissues at concentrations ranging from 0.2 to 0.6 μg g^-1^ in cereals and 0.6 to 1.2 μg g^-1^ in oilseeds crops (Haneklaus et al., 2015). Low concentrations of lanthanide application, above a physiological threshold, can be beneficial for plant growth through hormesis. Cereals and oilseed crops can tolerate rare earth element concentrations up to 1.0 and 3.0 μg g^-1^, respectively (Hu et al., 2002) This highlights that the range of lanthanide concentrations in foliar- or soil-applied fertilizers that can elicit beneficial effects on crop yields is small.

Two rare earth elements, cerium and lanthanum, have been previously associated with increased plant growth (Ippolito et al., 2011). For example, application of cerium nitrate or lanthanum nitrate at 0.5 to 10 μM increased primary root length, bolting, plant height, dry weight and average flower number of *Arabidopsis thaliana*; application of 0.5 μM of each cerium nitrate and lanthanum nitrate induced floral initiation and enhanced the effects of 10^-6^ M isopentenyl adenosine on root growth, plant height and flowering (He and Loh, 2000). In another study, addition of lanthanum or cerium, at 0.5 to 25 mg L^-1^, reduced primary root elongation, reduced root and shoot dry weight and reduced Ca, Mg, K, Cu and Zn concentrations in wheat seedlings (Hu et al., 2002).

Studies have demonstrated that rare earth metal-containing fertilizers can cause oxidative stress in plants which can have positive or negative effects on plant growth depending on application rate (Wang et al., 2007; Ippolito et al., 2011). For example, negative effects of lanthanides may result from changes in enzyme function, replacement of essential metals in pigments or production of reactive oxygen species (Babula et al., 2009).

Data collected in soybean showed that application of low concentrations of lanthanum stimulated photosynthetic rate and increased total chlorophyll content; these effects were associated with slightly increased root and shoot biomass (de Oliveira et al., 2015). In contrast, higher concentrations of lanthanum application reduced soybean growth. Stimulation of chlorophyll synthesis and enhanced root and shoot growth has also been reported in wheat, cucumber, soybean and corn plants (Wu et al., 1983; Wu et al., 1984).

### Rare earth element effects on plant nutrient uptake

Lanthanide-based fertilizer application has been reported to significantly alter nutrient concentration in roots and shoots of various crop plants under field and greenhouse conditions. For example, Chen and Zhao (2009) found that lanthanum application decreased Ca, Mg, Cu, Zn and Fe concentration in shoots and Cu and Zn concentrations in roots. In contrast, de Oliveira et al. (2015) found that lanthanum application increased Ca, P, K and Mn contents but reduced Cu and Fe contents of soybean plants. These effects have been hypothesized to be related to synergistic and antagonistic interactions of uptake between plant nutrients and lanthanides (Hu et al., 2004; Haneklaus et al., 2015). These effects may also result from improved Ca and Mn bioavailability in soil (Chang, 1991).

### Rare earth element fertilizer effects on crop yield

Previous experiments have shown variable effects of rare earth element fertilizers on crop yields (EI-Ramady, 2008). The main determinant of lanthanide fertilizers on crop yields depends on the concentration of lanthanides – benefits of these fertilizers are maximized when soil lanthanide concentration is below 10 mg kg^-1^ while benefits are unlikely from additional lanthanide fertilizer application when soil lanthanide concentration is above 20 mg kg^-1^ (Haneklaus et al., 2015). Average reported yield increases range from 5 to 15 % for crops including corn, cotton, potato, rice, soybean, sugar beet and wheat; the largest reported yield increases are up to 103 %. However, not all trials have reported crop yield increases which suggests that other factors, including crop, soil fertility, fertilizer application rate and/or environmental conditions also contribute to treatment effects in field trials.

## Materials and methods

### Fertilizer treatment description

REAP is a liquid rare earth and micronutrient supplement derived from potassium hydroxide, calcium sulfate, magnesium sulfate and ferric sulfate. The product contains lanthanum (derived from La^3+^), cerium (derived from Ce^3+^) and ethylenediamine tetraacetic acid (EDTA) as a chelate. Nutrient concentrations are given in Table 1.

**Table 1.** Nutrient contents of REAP.

| Elements | Concentration (%) |
| --- | --- |
| Available Phosphoric Acid ($\text{P}_2\text{O}_5$) | 1.00 |
| Soluble Potash ($\text{K}_2\text{O}$) | 0.30 |
| Calcium (Ca) | 0.04 |
| Magnesium (Mg) | 0.04 |
| Sulfur (S) | 0.10 |
| Iron (Fe) | 0.02 |
| Lanthanum (La) | 0.02 |
| Cerium (Ce) | 0.05 |

### Experimental design

Two experiments were conducted to determine the effects of REAP on yield and physiological performance of greenhouse crops. In the first experiment (conducted in September to November 2004), REAP was applied as a supplemental fertilizer at rates of 100, 250 or 500 ppm (equivalent to 0.378, 0.946 and 1.89 mL gallon^-1^, respectively); in the second experiment (conducted in January to September 2005), REAP was applied as a supplemental fertilizer at rates of 500, 1000, or 2000 ppm (equivalent to 1.89, 3.79 and 7.57 mL gallon^-1^, respectively). In both experiments, the control did not receive any supplemental fertilizer. Experiments used a randomized complete block design with three blocks to account for variability in light levels and temperature in the greenhouse. In total, the experiments used four crops: lettuce, pepper, tomato and cantaloupe. Crop varieties and number of plants per experiment are shown in Table 2.

**Table 2.** Crop varieties and number of replicates per treatment for each experiment.

| **Crop** | **Variety** | **Experiment 1** |  | **Experiment 2** |
| --- | --- | --- | --- | --- |
| Lettuce | Romaine Triton | 16 | 16 | 12 |
|  | Green Salad Bowl | 16 | 12 | - |
|  | Red Sails | - | - | 12 |
| Pepper | California Wonder | 6 | 18 | 12 |
|  | New Ace | 6 | 18 | - |
| Tomato | Bush Early Girl | 6 | - | - |
|  | Red Husky | - | - | 12 |
| Cantaloupe | - | - | - | 30 |

### Measurements and harvest

For pepper plants, samples of were taken to measure biomass production in Experiment 1 and stem diameter and biomass production were recorded in Experiment 2. For tomato plants, gas exchange measurements (photosynthesis rate, transpiration rate and stomatal conductance) were taken in Experiment 2. For cantaloupe, the weight, number of plants and weight per plant were recorded in Experiment 2. All crops were harvested once they reached physiological maturity and yield was recorded. For tomato plants, nutrient analysis was conducted to determine total N, total P, K, Ca, Mg, S, and Na (all in %) and Fe, Zn, Mn, Cu, and B (all in mg kg^-1^). Roots were harvested and photographer for Romaine Triton (lettuce) and California Wonder (pepper) in Experiment 1 and for California Wonder (pepper) and Red Husky (tomato) in Experiment 2.

### Statistical analysis

Data were analyzed using two-way ANOVA in Excel. One factor was treatment and the second factor was the block effect. When models were statistically significant (probability of F < 0.05) or showed a trend effect (probability of 0.05 < F < 0.10), differences between treatments were calculated using Fisher’s least significant difference. Results are reported as significant when treatment differences have p < 0.05 and are reported as a trend when p < 0.10. Differences that are not statistically significant but result in a 10 % or greater change to a plant physiological parameter with REAP application are also mentioned.

## Results

### REAP increases crop yield

In the first experiment, REAP did not have a significant effect on yields for any crops at any of the rates tested (Figures 1-3). However, pepper biomass production was increased by 17.2 % when REAP was applied at 500 ppm compared to the control treatment (Figure 2A). Root biomass of Romaine Triton and California Wonder appeared to increase with REAP application at 500 ppm based on visual assessment (Figure 1E, F and Figure 2E, F) but the magnitude of this increase was not quantified. However, in the second experiment, REAP lead to significant yield increases for lettuce, pepper, tomato and cantaloupe (Figures 4-7). Romaine Triton yield tended (p < 0.10) to be increased by 19.0 % when REAP was applied at 1000 ppm (Figure 4A). Red Sails lettuce yield increased by 13.5 and 16.1 % compared to the control for the 1000 and 2000 ppm REAP treatments, respectively (p < 0.05 for both) (Figure 4B). The increases in lettuce yield appeared to be associated with increased biomass per head of lettuce (+10.6 % when REAP was applied at 1000 ppm for Romaine Triton and + 12.6 % when REAP was applied at 2000 ppm for Red Sails), however these effects were not statistically significant (Figure 4C, D). Pepper yield was increased by 11.4, 44.2 and 31.9 % when REAP was applied at 500, 1000 and 2000 ppm, however only the increased associated with a REAP application of 1000 ppm tended (p < 0.10) to be significant (Figures 5A). Similarly, pepper stem diameter increased by 12.4 and 11.0 % when REAP was applied at 1000 and 2000 ppm (Figure 5B), respectively; pepper biomass production was increased by 16.7, 36.1 and 39.7 % when REAP was applied at 500, 1000 and 2000 ppm, respectively (Figure 5C). The root biomass of California Wonder and New Ace appeared to increase with REAP application at 500 ppm based on visual assessment (Figure 5D, E), but the magnitude of this increase was not quantified. Increased yield appeared to be associated with increases in the number and yield of marketable vs. culled peppers over the study period and an increased rate of yield accumulation (Figure 5F-J). Tomato yield was increased by 13.5 and 16.1 % when REAP was applied at 1000 and 2000 ppm, respectively (p < 0.05 for both) (Figure 6A). Again, the root biomass of Red Husky appeared to increase with REAP application based on visual assessment (Figure 6B), but the magnitude of this increase was not quantified. This appeared to be associated with an increase in the number of marketable tomatoes and a reduction in the number of culled tomatoes and an increased rate of tomato yield accumulation when REAP was applied (Figure 6C-G). Finally, cantaloupe yield was increased by 30.0, 73.7 and 41.3 % when REAP was applied at 500, 1000 and 2000 ppm, respectively (p < 0.10, p < 0.05 and p < 0.05, respectively) (Figure 7A). This appeared to be associated with an increase in the number of melons per plant and mass per melon (Figure 7B, C). Root biomass of cantaloupe did not appear to respond as strongly to REAP application compared to the other crops (Figure 7D).

**Figure 1:**
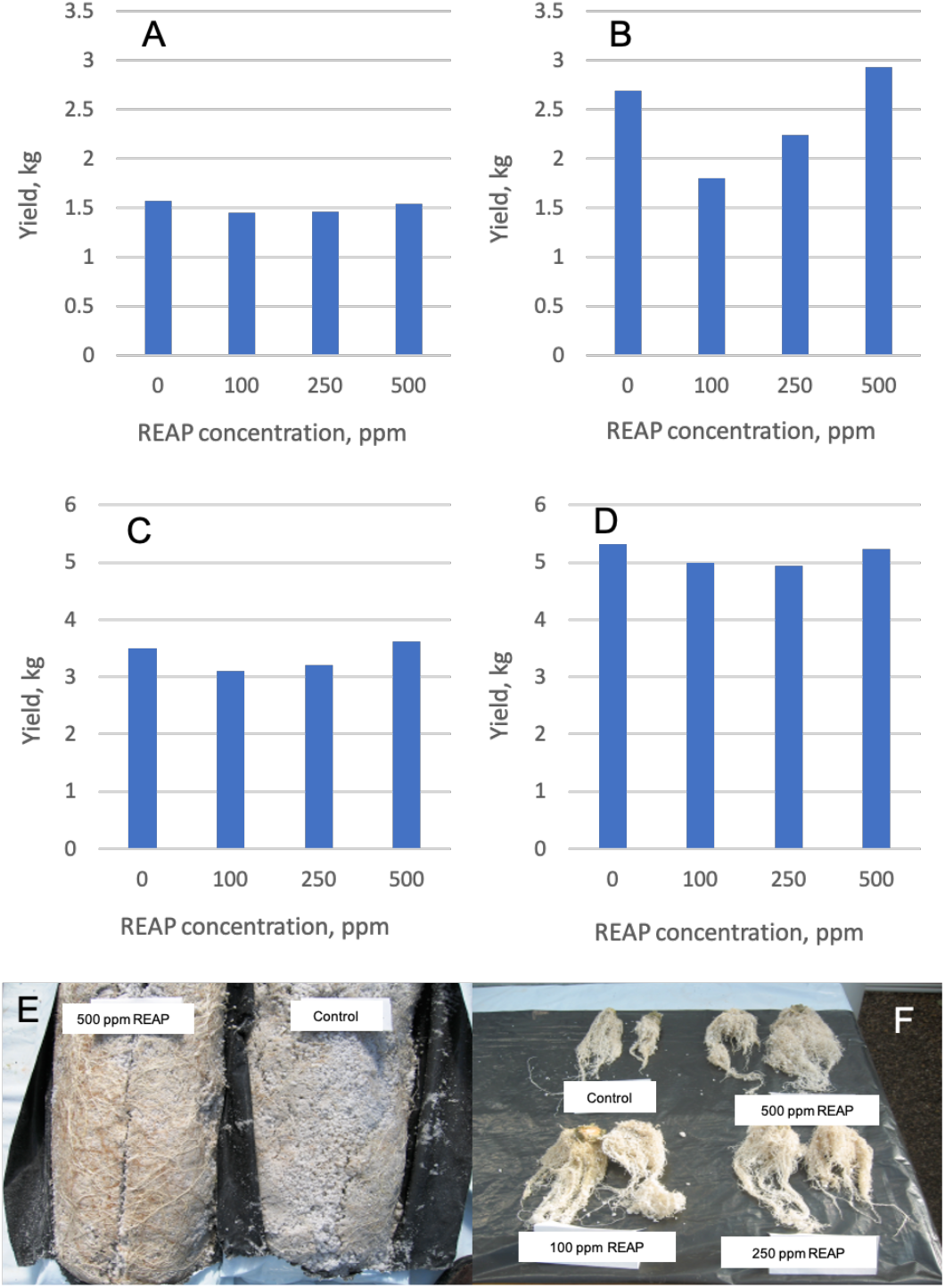
Effects of REAP on lettuce yield and root biomass in Experiment 1. Yield for Green Lettuce Bowl in run 1 (A) and run 2 (B). Yield for Romaine Triton in run 1 (C) and run 2 (D). Romaine Triton roots (E, F).

**Figure 2:**
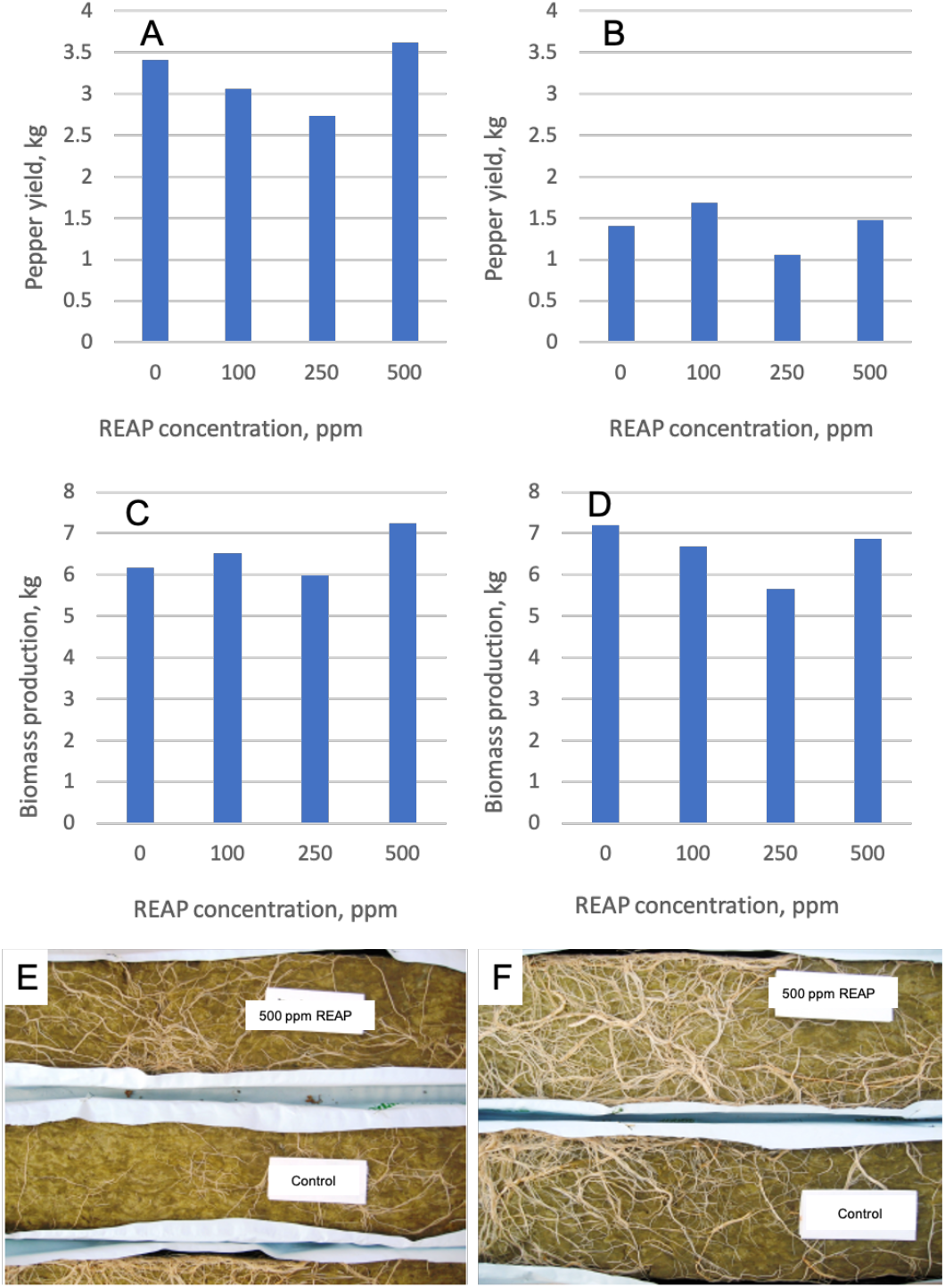
Effects of REAP on pepper yield, biomass production and root biomass in Experiment 1. Yield for New Ace (A) and California Wonder (B). Biomass production for New Ace (C) and California Wonder (D). Roots of New Ace (E) and California Wonder (F).

**Figure 3:**
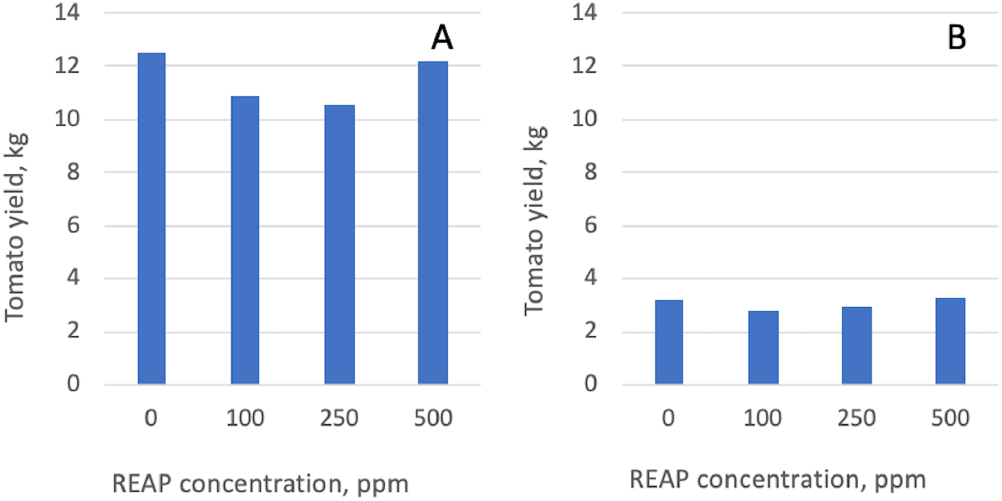
Effects of REAP on tomato yield in experiment 1. Yield for Bush Early Girl (A) and Patio Hybrid (B).

**Figure 4:**
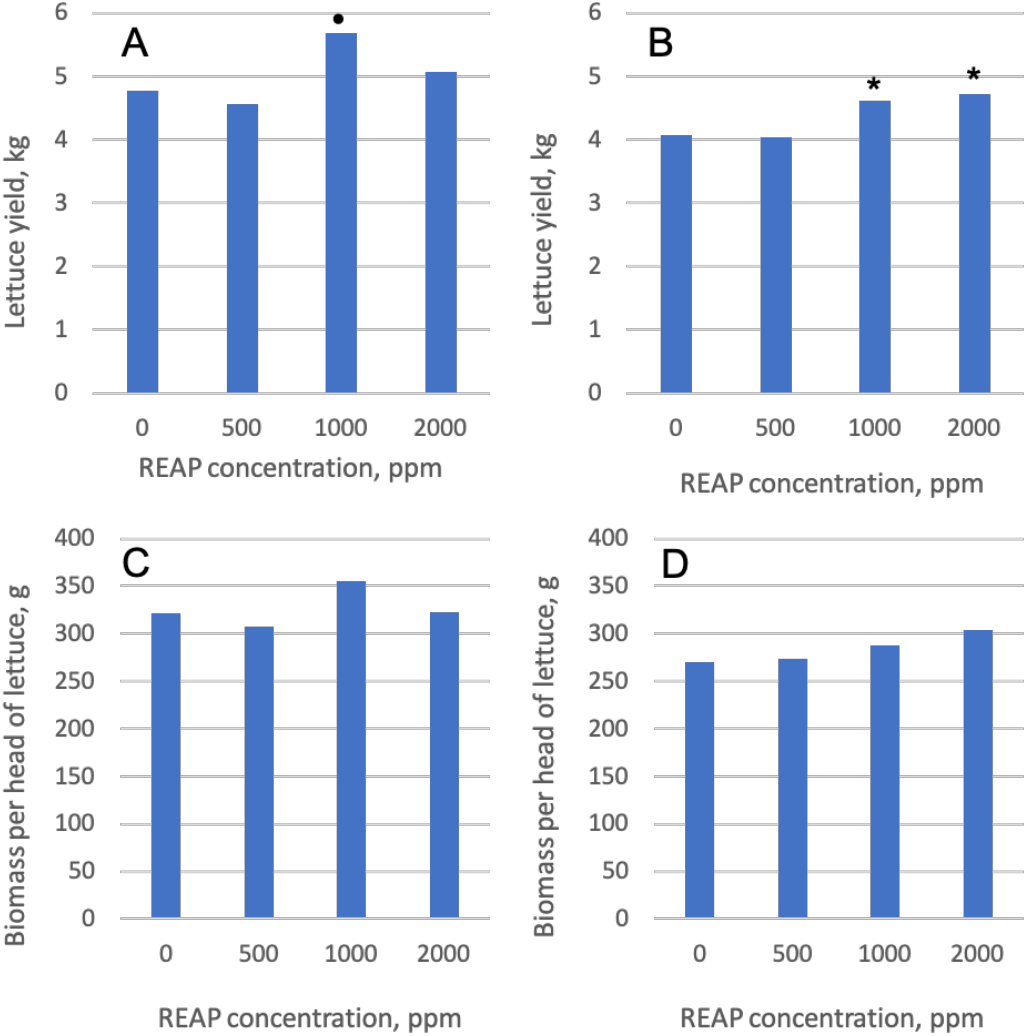
Effects of REAP on lettuce yield and biomass per head of lettuce in Experiment 2. Yield for Romaine Triton (A) and Red Sails (B). LSD for Red Sails = 0.440 (p < 0.05), 0.497 (p < 0.10); LSD for Romaine Triton = 0.551 (p < 0.05), 0.408 (p < 0.10); • tended to increase yield at p < 0.10; * increased yield at p < 0.05. Biomass per head of lettuce for Romaine Triton (C) and Red Sails (D).

**Figure 5:**
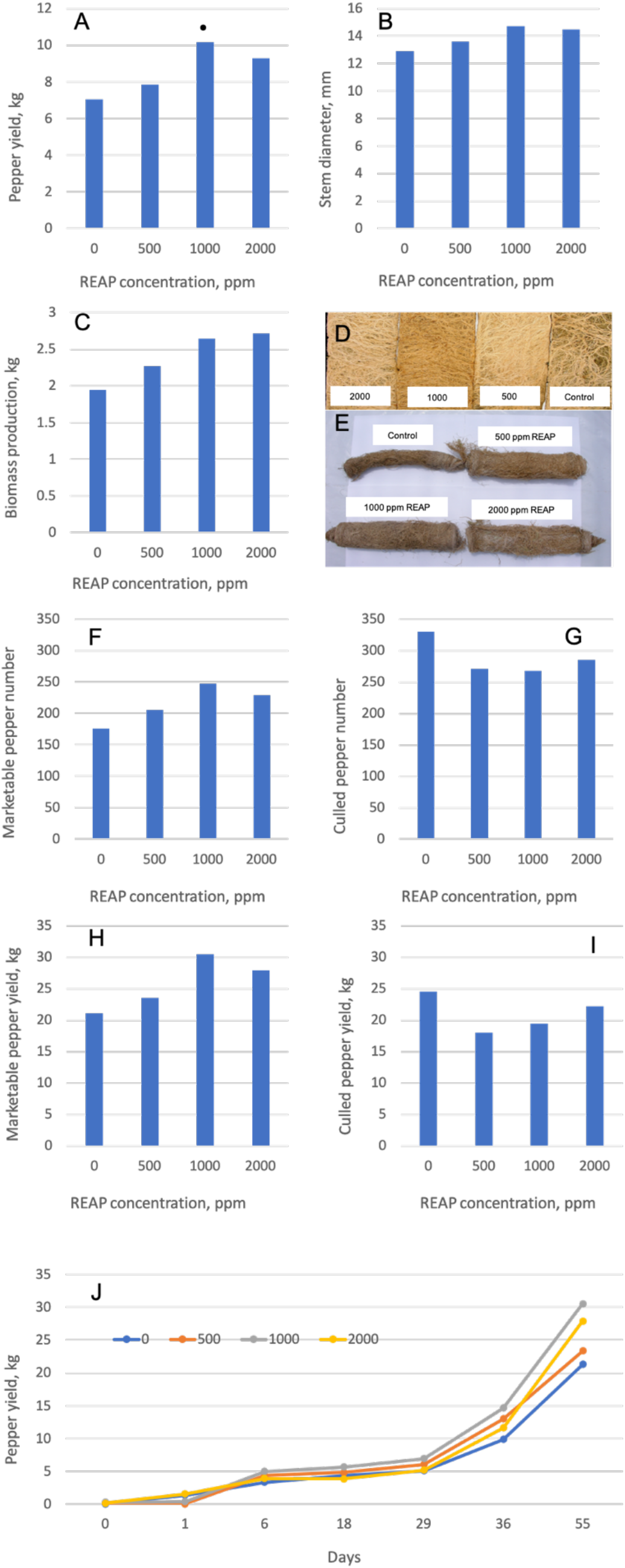
Effects of REAP on pepper (California Wonder) in Experiment 2. California Wonder yield (A), stem diameter (B), biomass yield (C), root biomass (D) and marketable vs. culled pepper number (F, G) and yield (H, I). For (A), LSD = 3.984 (p < 0.05), 2.947 (p < 0.10), • tended to increase yield at p < 0.10. For (F-I), marketable and culled pepper number and yield are for the period from May 31 to July 26.

**Figure 6:**
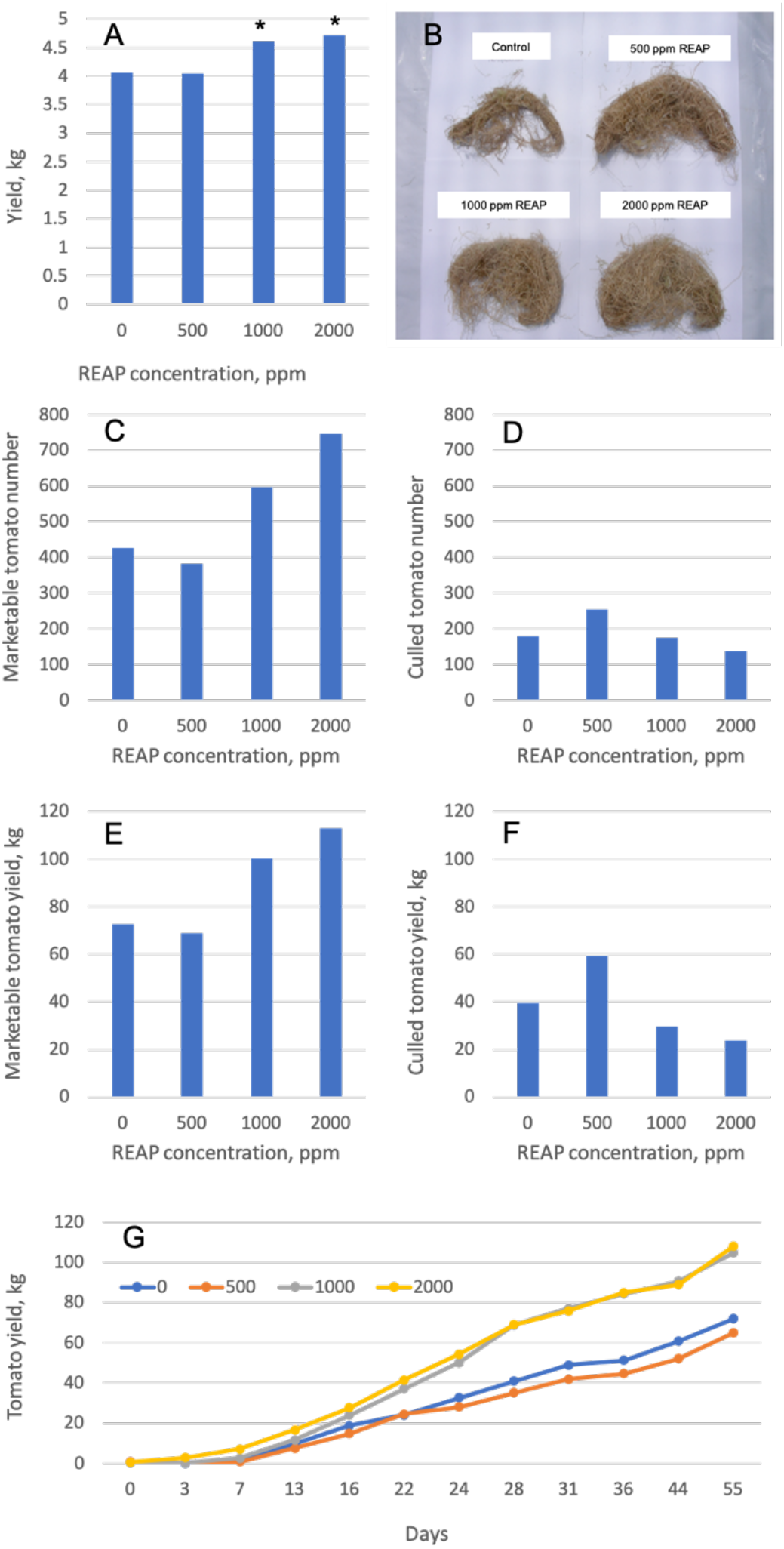
Effects of REAP on tomato (Red Husky) in Experiment 2. Red Husky yield (A), root biomass (B), marketable vs. culled tomato number (C, D) and yield (E, F) and cumulative tomato yield (G). For (A), LSD = 16.093 (p < 0.05), 11.901 (p < 0.10), * increased yield at p < 0.05. For (C-F), marketable and culled tomato number and yield are for the period from May 24 to July 17.

**Figure 7:**
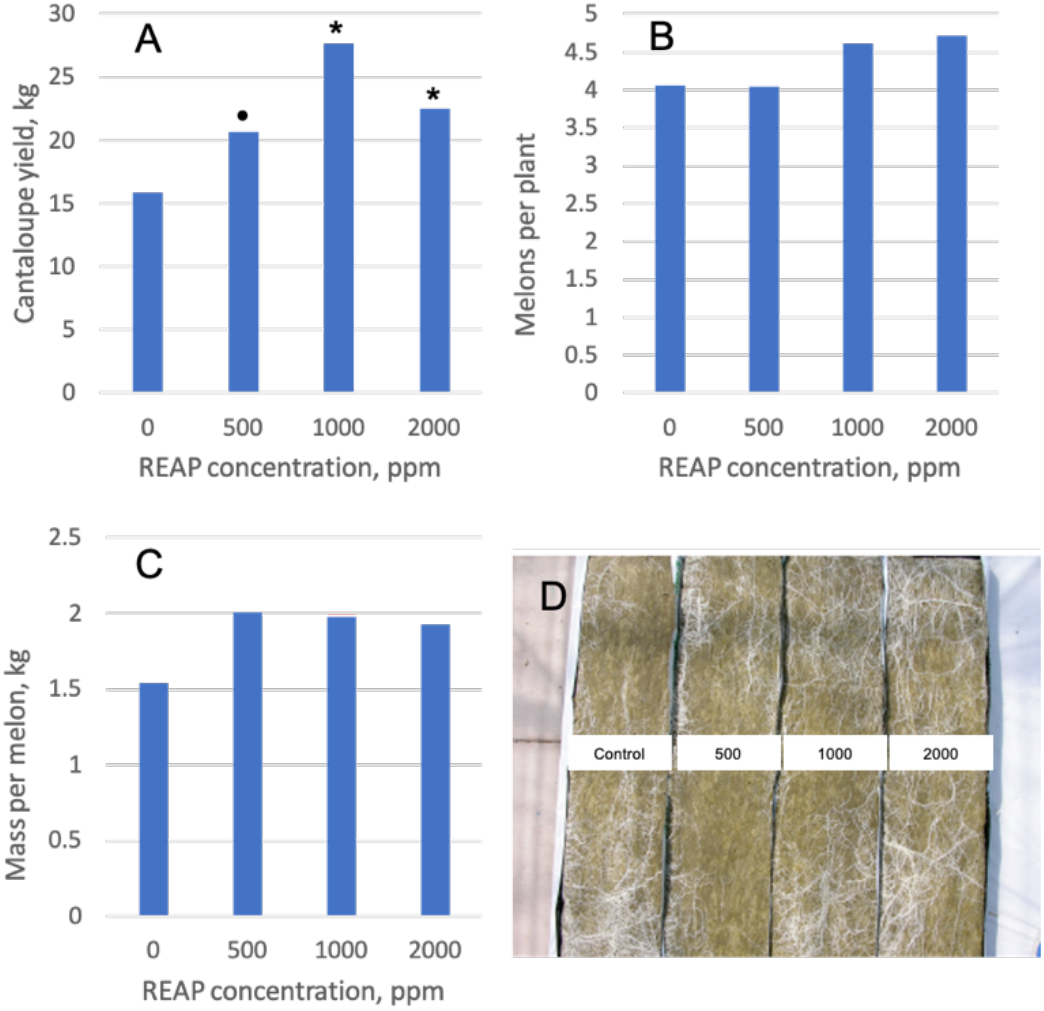
Effects of REAP on cantaloupe in Experiment 2. Cantaloupe yield (A), number of melons per plant (B), mass of melon per plant (C) and root biomass (D). For (A), LSD = 6.127 (p < 0.05), 2.353 (p < 0.10), • tended to increase yield at p < 0.10, * increased yield at p < 0.05.

### REAP effects on plant physiology, nutrient content and root biomass

REAP did not alter gas exchange measurements (photosynthesis rate, transpiration rate, or stomatal conductance) of tomato plants in Experiment 2 on either date when measurements were taken (Table 3). While REAP has some small effects on tomato nutrient analysis, differences between treatments were not statistically significant (Table 4). For example, total P concentration was increased by 13.4, 26.9 and 41.8 % compared to the control when REAP was applied at 500, 1000 or 2000 ppm, respectively; Zn concentration was increased by 38.5, 43.6 and 74.4 % compared to the control when REAP was applied at 500, 1000 or 2000 ppm, respectively; Cu increased by 40 % compared to the control when REAP was applied at 2000 ppm. Ca, Mg, Na, Fe were 20.8, 15.6, 41.2 and 27.0 % lower than the control, respectively, when REAP was applied at 2000 ppm.

**Table 3.** Gas exchange measurements of Experiment 2 tomato plants taken on two dates.

| Hy Yield<br>rate,<br>ppm | 01/24/2005 |  | 06/02/2005 |  |  |
| --- | --- | --- | --- | --- | --- |
| | Photosynthesis<br>rate,<br>$\mu\text{mol m}^{-2} \text{s}^{-1}$ | Transpiration<br>rate,<br>$\text{mmol m}^{-2} \text{s}^{-1}$ | Photosynthesis<br>rate,<br>$\mu\text{mol m}^{-2} \text{s}^{-1}$ | Transpiration<br>rate,<br>$\text{mmol m}^{-2} \text{s}^{-1}$ | Stomatal<br>conductance,<br>$\mu\text{mol m}^{-2} \text{s}^{-1}$ |
| 0 | 14.79 | 6.56 | 17.23 | 5.44 | 534 |
| 500 | 14.90 | 6.69 | 18.58 | 5.69 | 571 |
| 1000 | 14.03 | 6.67 | 18.32 | 5.68 | 572 |
| 2000 | 14.53 | 6.87 | 17.17 | 5.65 | 563 |

**Table 4.** Tomato nutrient analysis (Experiment 2).

| Nutrient | Hy Yield application rate, ppm |  |  |  |
| --- | --- | --- | --- | --- |
|  | 0 | 500 | 1000 | 2000 |
| Total N (%) | 3.6 | 3.8 | 3.5 | 3.6 |
| Total P (%) | 0.67 | 0.76 | 0.85 | 0.95 |
| K (%) | 11.0 | 11.0 | 11.0 | 11.0 |
| Ca (%) | 2.40 | 2.50 | 2.10 | 1.90 |
| Mg (%) | 0.64 | 0.65 | 0.60 | 0.54 |
| S (%) | 0.34 | 0.30 | 0.34 | 0.38 |
| Na (%) | 0.17 | 0.16 | 0.13 | 0.1 |
| Fe (mg kg<sup>-1</sup>) | 37 | 27 | 29 | 27 |
| Zn (mg kg<sup>-1</sup>) | 39 | 54 | 56 | 68 |
| Mn (mg kg<sup>-1</sup>) | 61 | 62 | 61 | 67 |
| Cu (mg kg<sup>-1</sup>) | 5 | 5 | 5 | 7 |
| B (mg kg<sup>-1</sup>) | 27 | 26 | 27 | 27 |

## Discussion

### REAP effects on crop yield

The results of this experiment are in line with previous reports of lanthanide fertilizers on crop yield. For example, we found that at lower application rates, in Experiment 1, the lanthanide-based fertilizer did not alter crop yield. Previous data from the literature shows that rare earth element fertilizers must be applied at a minimum threshold concentration to achieve beneficial effects on crop yield (Haneklaus et al., 2015). Accordingly, at higher application rates used in Experiment 2, we found that REAP improved yield of lettuce, pepper, tomato and cantaloupe. This finding supports the hypothesis that lanthanides provide a hormetic effect on plant growth – there is a specific, relatively low dose, of lanthanide concentration that elicits positive effects on crop growth. In this experiment, evidence of lanthanide toxicity was not observed, even at the highest application rate of 2000 ppm. However, previous research has shown that at higher application rates, lanthanide-based fertilizers can reduce crop yield. Future experiments could examine the highest rate of REAP that is associated with positive or neutral effects on crop yield for various crops. Furthermore, it is likely that different crops have differing levels of lanthanide concentration and maximum application rates should be tested for each crop.

In this experiment, REAP did not have significant effects on nutrient concentration or gas exchange rates in tomato plants. Previous research demonstrated that when Mg supply was low in soil, La and Ce could replace Mg to improve photosynthetic rate (Hong et al., 2002). However, in the current experiment, plants received fertigation at near-optimum rates so this mechanism of growth promotion may not have been relevant. Instead, the yield responses observed in this study may be associated with other mechanisms of lanthanide-associated growth stimulation such as interactions with abscisic acid to alter hydrogen peroxide production (Wang et al., 2014). Future experiments could examine plant hormone and reactive oxygen species levels in response to lanthanide-containing fertilizers to better understand the mode-of-action.

Application of REAP fertilizer appeared to increase root biomass accumulation in this study. Future experiments could use root architecture analysis, using CT or flat-bed scanning (e.g., WinRhizo), to quantify the changes that occur as a result of lanthanide fertilizer application. Previously, increased tuber yield of *Pseudostellaria heterphylla* was associated with reduced midday depression of gas exchange measurements (Ma et al., 2017).

Data from the literature suggests that crop yield enhancements associated with lanthanide-based fertilizers may result from improved plant stress responses and reduced salinity and heavy metal uptake. For example, when 0.1 mM lanthanum nitrate was applied to *Saussurea involucrata*, a plant used in traditional Chinese medicine, growth inhibition associated with salinity (NaCl) stress was reduced, leaf water potential, water content, and proline and protein contents were increased, while malondialdehyde contents were decreased (Xu et al., 2007). In the same study, lanthanum application was also associated with increased antioxidant enzyme activities, reduced decomposition of photosynthetic pigmentation. Similar effects of lanthanum on salinity stress have been reported in other crops, including *Vigna radiata* (Shan and Zhao, 2014), tomato (Huang and Shan, 2018) and corn (Liu et al., 2016) while cerium has also been reported to reduce salinity stress in corn (Hu and Shan, 2018). Lanthanum has also been reported to reduce the impact of cadmium stress by regulating plant ascorbate and glutathione metabolism and simultaneously reducing cadmium uptake in corn plants (Dai et al., 2017). Research has also demonstrated that lanthanum reduces both uptake and translocation of cadmium in wheat plants (Yang et al., 2019). In addition, lanthanum can also reduce the impact of chromium stress in corn plants through similar mechanisms (Dai and Shan, 2019). Improved plant resistance to salinity stress and reduced uptake of heavy metal contaminants is particularly relevant for hydroponic crops where salinity stress can reduce crop yields while heavy metal contamination is undesirable for crops destined for human consumption.

Future experiments could measure salinity, nutrient, heavy metal and lanthanide uptake in tissues of plants receiving REAP fertilizer supplementation in relation to its effects on root architecture and biomass, and on translocation within the plant. Lanthanide fertilizers may also reduce heavy metal and/or lanthanide contaminants levels in aerial plant tissue by sequestration of these compounds in root tissues (Haneklaus et al., 2015) which should be evaluated carefully in future work. Previous data suggests that fertigation application of lanthanide fertilizers to crops where aboveground plant portions are consumed, such as those considered in this study, are less likely to accumulate lanthanides in edible plant parts. For example, when rare earth element fertilizer was applied to corn, at less than 10 kg ha^-1^, these elements did not accumulate in aboveground biomass (Haneklaus et al., 2015). Previous experiments have shown that altered root architecture and biomass can reduce fertilizer requirements (Backer et al., 2017; d’Aquino et al., 2009; Ramos et al., 2016) which suggests that root architecture changes associated with REAP application may lead to reduced fertilizer requirements while maintaining crop yields.

In addition, future experiments could examine the effects of lanthanide-based fertilizers on post-harvest crop quality parameters and nutritional content since these fertilizers can alter plant nutrient uptake and antioxidant enzyme system (Ramos et al., 2016). Lanthanides have previously been shown to alter plant secondary metabolite concentrations; often the accumulation of these molecules alters crop nutritional value and/or flavour profile (Ma et al., 2014). For example, in tomato cultivation, lanthanide fertilizers may increase carotenoid compounds such as *α*- and *β*-carotene, lutein and lycopene and vitamin C concentration, while in cannabis cultivation this may have desirable effects on crop potency (i.e., cannabinoid concentration in flowers) or aroma (i.e., terpenoid or flavonoid profiles in flowers) for plants growing in controlled environments (data not shown). Similarly as for the effects of rare earth elements on crop yield, the effects on the biosynthesis of plant secondary metabolites depends on the application rate. For example, the concentration of vitamin C in strawberry increased at low concentrations of lanthanum nitrate application but was reduced at higher application rates (Shan et al., 2017). There is value in determining the appropriate application rate of rare earth element fertilizers to impact plant secondary metabolite biosynthesis since these changes are important for consumers and improve marketability of the crop.

## Conclusions

Greenhouse experiments that REAP increased greenhouse crop (lettuce, pepper, tomato and cantaloupe) yield when used as a supplemental fertilizer in a constant liquid feed system at application rates ranging from 500 to 1000 ppm. This appeared to be the result of the hormetic effects previously reported for lanthanide-based fertilizers since lower application levels (100 to 250 ppm) to not affect yield of these crops. Yield increases were associated with apparent increased biomass production for pepper and tomato; however, this increase was not quantified. Future studies could examine how root architecture and biomass accumulation respond to supplemental lanthanide fertilizer application and whether this is associated with increased nutrient uptake and/or improved nutrient concentration balance in plant tissue. In this study, tomato nutrient concentration and gas exchange measurements were not significantly impacted by supplemental lanthanide fertilizer application. Future studies could confirm this effect and examine other possible mechanisms underlying growth promotion such as activity of oxidative-stress-related enzymes in plant tissue. Finally, future studies could consider the impact of supplemental lanthanide fertilizers on crop nutritional and/or quality parameters that may be associated with altered stress response physiology.

## Acknowledgements

We are grateful to the technicians who collected the data in this manuscript and to the anonymous reviewers for their constructive feedback.

